# The Network Zoo: a multilingual package for the inference and analysis of biological networks

**DOI:** 10.1101/2022.05.30.494077

**Authors:** Marouen Ben Guebila, Tian Wang, Camila M. Lopes-Ramos, Viola Fanfani, Deborah Weighill, Rebekka Burkholz, Daniel Schlauch, Joseph N. Paulson, Michael Altenbuchinger, Abhijeet Sonanwane, James Lim, Genis Calderer, David van Ijzendoorn, Daniel Morgan, Alessandro Marin, Cho-Yi Chen, Alex Song, Kate Shutta, Dawn DeMeo, Megha Padi, John Platig, Marieke L. Kuijjer, Kimberly Glass, John Quackenbush

## Abstract

Inference and analysis of cellular biological networks requires software tools that integrate multi-omic data from various sources. The Network Zoo (netZoo; netzoo.github.io) is an open-source software suite to model biological networks, including context-specific gene regulatory networks and multi-omics partial correlation networks, to conduct differential analyses, estimate community structure, and model the transitions between biological states. The netZoo builds on our ongoing development of network methods, harmonizing the implementations in various computing languages (R, Python, MATLAB, and C) and between methods to allow a better integration of these tools into analytical pipelines. To demonstrate the value of this integrated toolkit, we analyzed the multi-omic data from the Cancer Cell Line Encyclopedia (CCLE) by inferring gene regulatory networks for each cancer cell line and associating network features with other phenotypic attributes such as drug sensitivity. This allowed us to identify transcription factors that play a critical role in both drug resistance and cancer development in melanoma. We also used netZoo to build a pan-cancer, multi-tiered CCLE map and used it to identify known metabolic hallmarks of cancer and to estimate novel context-specific elements that mediate post-transcriptional regulation. Because the netZoo tools are open-source and there is a growing community of both users and developers, we built an ecosystem to support community contributions, share use cases, and visualize networks online. As additional data types become available and our suite of methods grows, we will expand “the zoo” to incorporate an increasingly sophisticated collection of tools for network inference and analysis.

## Background

Biological phenotypes are driven by a complex network of interacting elements that defines cell types and determines response to perturbations [1]. Defining these interactions can be modeled by assessing physical binding between biological elements [2], their co-expression [3], and their co-dependency [4] to identify functional modules that together control the emergence of a given phenotype. A particular type of network are gene regulatory networks (GRNs) that are comprised of regulators and their target genes. One type of regulators are transcription factors (TFs), regulatory proteins that bind to DNA to activate or repress gene transcription. TFs often form complexes that act together to regulate transcription [5–7] and TF activity can be further influenced by epigenetic modifications such as promoter methylation or histone acetylation [8]. Other regulators of gene expression include microRNAs (miRNAs) that act post-transcriptionally, primarily to degrade and subsequently repress the expression of their mRNA target [9, 10]. These and other factors together modulate the expression of the more than twenty-five thousand protein-coding genes in the genome, altering cellular processes and giving cells the potential to respond to various stimuli [7].

Despite rapid advances in sequencing technologies, the size and complexity of biological networks put them out of reach of direct measurement [6]. Consequently, there have been many attempts to model and represent biological networks using computational methods [3, 6, 11–13], although not all model gene regulatory processes.

Our group has developed a number of robust methods for GRN inference and analysis, each of which takes advantage of multiple data types available in individual studies. Each method is based on modeling known biological interactions and seeks consistency between a variety of input data sources. Our methods for reconstructing networks include PANDA [14] and OTTER [15] for modeling TF-gene regulatory processes, DRAGON [16] for estimation of multi-omic networks based on GGMs, and PUMA [17] which adds miRNA-gene post-transcriptional regulation to TF-gene interactions In addition, SPIDER [18] reconstructs networks by accounting for chromatin state, EGRET [19] include genotype information in network inference, and LIONESS [20] estimates network models for individual samples. Another set of methods has been developed for network analysis including CONDOR [21] for modeling and detection of communities in expression quantitative trait locus (eQTL) networks, ALPACA [22] and CRANE [23] for identifying communities within networks and how communities change between states, and MONSTER [24] to estimate TFs that drive the transition between network states. SAMBAR [25] allows us to group biological samples based on how genetic variants alter functional pathways, and, finally YARN [26] is a tissue-aware implementation of smooth quantile normalization for multi-tissue gene expression data.

Many of these methods share a methodological and philosophical framework that derives from the “No Free Lunch Theorem”—modeling of complex systems can be improved by incorporating domain-specific knowledge [27]. Also, many of them use an overlapping set of standard input data types and provide complementary views of biological networks. As such, they have often been used together. To facilitate their use and integration into analytical pipelines, we gathered these into the Network Zoo (netZoo; netzoo.github.io), a platform that harmonizes the codebase for these tools and provides implementations in R, Python, MATLAB, and C. In building netZoo, we also created the Zookeeper, a tool that helps ensure consistency of the codebase as it is continuously updated in response to user feedback. The netZoo serves as a centralized resource for biological networks including GRN inference and analysis by providing an ecosystem of tools available to both scientists and method developers and includes resources to integrate contributions, to share use cases [28], and to host and visualize networks [29].

To demonstrate the power of this unified platform, we used netZoo tools to build the first comprehensive collection of genome-scale GRNs for the cell lines in the Cancer Cell Line Encyclopedia (CCLE) [30–32]. We also used PANDA, LIONESS, and MONSTER to infer TF-gene targeting in melanoma to explore how regulatory changes affect disease phenotype, and used DRAGON to integrate nine types of genomic information and find multi-omic markers that are associated with drug sensitivity.

## Results and discussion

### netZoo integrates network inference and downstream analyses

Regulatory processes drive gene expression and help define both phenotype and the ability of a biological system to respond to perturbations. However, identifying context-specific regulatory processes is difficult because the underlying regulatory network is often unobserved [6]. Several netZoo methods address this challenge by integrating multiple sources of available data to infer TF-gene regulation. PANDA [14] builds a regulatory TF-gene network by first positing a prior regulatory network and then iteratively optimizing its structure by seeking consistency between gene co-expression and TF protein-protein interactions (PPIs). The prior regulatory network can be constructed by scanning the sequence of the promoter region of target genes (for example, by using FIMO [33]) for transcription factor binding sites (TFBS) using TF motifs taken from catalogs (such as CIS-BP [5]). The input TF PPI data can be obtained from resources such as STRING [2] and gene co-expression is obtained from the particular experiment being analyzed. The inference is based on the concept that interacting TFs co-regulate their target genes and co-expressed genes are potentially regulated by the same sets of TFs. PANDA uses message passing to iteratively update all three data sets, maximizing consistency between them, until it converges on a data set-specific regulatory network with interaction scores between TFs and their regulated targets.

Other methods in netZoo were built using PANDA’s conceptual framework. OTTER [15] takes the same input but uses regression as an alternative implementation of the network optimization solution. SPIDER [18] uses epigenetic data such as DNase-Seq measurements of DNA accessibility to inform the PANDA prior network on context-specific accessible chromatin regions. EGRET [19] uses cis-eQTL data to seed the method with genotype-specific priors. PUMA [17] extends PANDA’s regulatory framework by including miRNA target predictions in the initial prior network to capture both TF and miRNA regulation of target genes/mRNAs.

LIONESS [20] is a general-purpose method for single-sample network estimation that can be used with any network inference approach. It iteratively leaves out individual samples and uses linear interpolation to estimate sample-specific networks for each sample in the original sample set. LIONESS outputs individual sample edge weights which can be treated as inferred measures on each sample, allowing statistical comparisons to be performed on the associated networks. A key use case of LIONESS is to estimate sample-specific GRNs using PANDA.

DRAGON [16]is a flexible method for integrating multiple data sources into a GGM. GGMs differ from correlation networks in that partial correlation corrects for spurious correlations between modeled variables; the multi-omic networks estimated by DRAGON therefore represent direct associations between the different omic types included in the model. DRAGON differs from PANDA and similar Zoo tools for network estimation in that it estimates an undirected unipartite network rather than a bipartite GRN.

A second group of tools in netZoo were developed to identify and explore higher-order structure in biological networks [34, 35] by identifying highly connected network “communities” and comparing the structure of these communities between phenotypic states. CONDOR [21] identifies communities in bipartite graphs (including eQTL networks and GRNs), while ALPACA [22] finds differential community structures between two networks, such as in a case versus control setting, by going beyond the simple difference of edge weights and using the complete network structure to find differential communities. CRANE [23] extends ALPACA’s differential community estimation by assessing significance of differential modules and comparing to a baseline of network ensembles that were generated by preserving the specific structure and constraints of GRNs. An important use case of CRANE is modeling the transition between an initial and a final condition such as between healthy and disease states. A fourth method, MONSTER [24], treats the transition between related biological states as one in which a first network is subject to a regulatory transition that involves altering transcription factor connections to their target genes. Mathematically, MONSTER models such changes by identifying a “transition matrix” that maps an initial state network to a final state network to identify the TFs that have the largest effect on the structure of the network and therefore are likely to help drive the phenotypic transition.

Many of the netZoo tools share common methodological and computational cores and over the years we have used combinations of these tools to explore the regulatory features driving biological states [36, 37]. Harmonizing the implementation of these tools to create a unified resource, netZoo, facilitates interoperability and the seamless integration in a pipeline that connects network inference with downstream analyses (Figure 1) to generate hypotheses and actionable biological insights.

**Figure 1.**
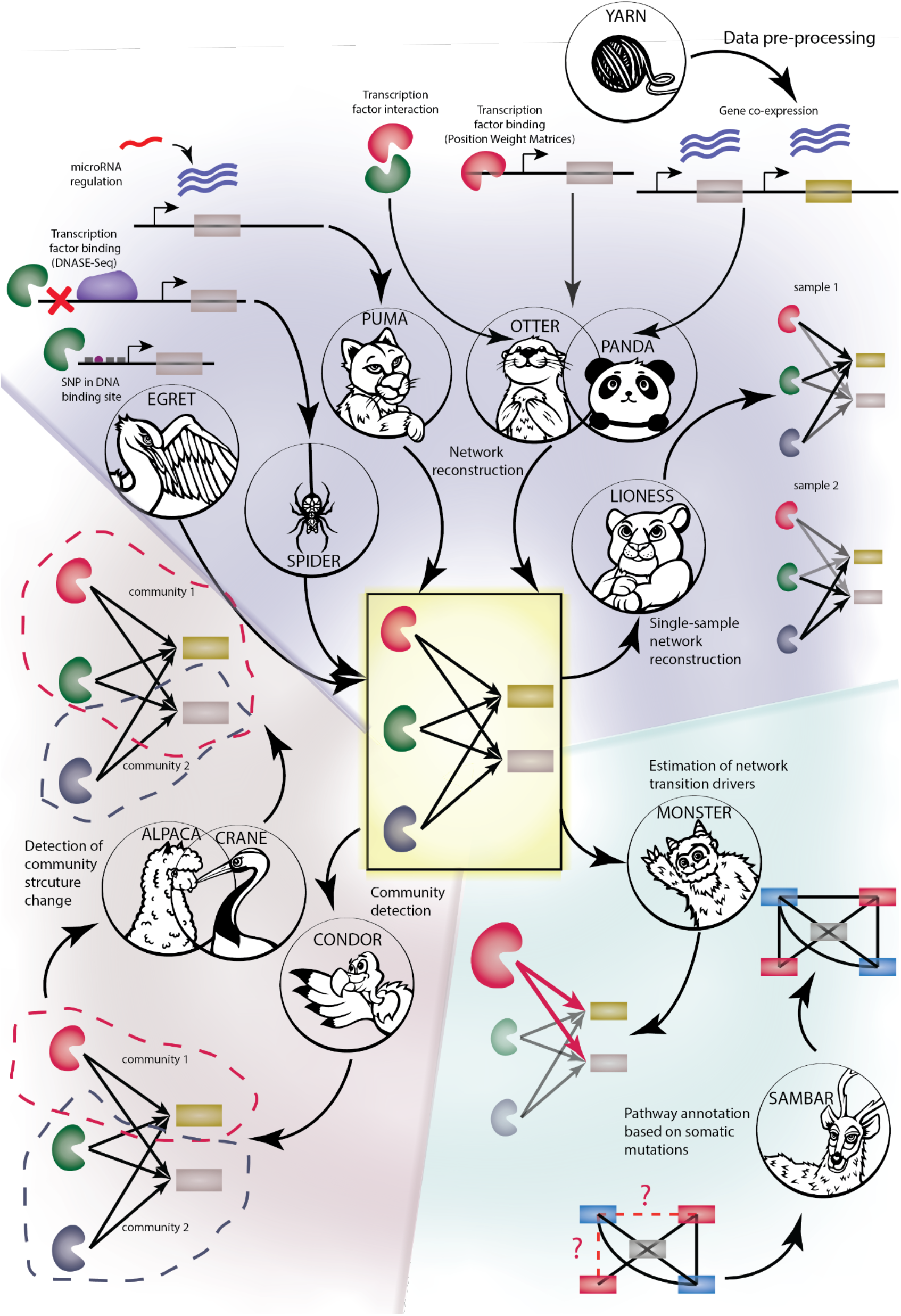
Inference and analysis of GRNs using netZoo. YARN allows us to normalize gene expression data to account for differences between tissues. Then, a first group of tools uses normalized gene expression data to infer cellular biological networks with a first set that reconstructs GRNs using multiple genomic modalities (PANDA, PUMA, OTTER, LIONESS, SPIDER, EGRET) and a second method for multi-omic correlation-based networks (DRAGON). A second group (CONDOR, ALPACA, CRANE) identifies communities in the networks and finds differential community structures between two networks of interest. Finally, MONSTER estimates a transition matrix between two networks representing an initial and a final state, and SAMBAR de-sparsifies mutation data using biological pathways. The methods represented in this figure are YARN, PANDA, PUMA, OTTER, LIONESS, SPIDER, EGRET, CONDOR, ALPACA, CRANE, MONSTER, and SAMBAR. SNP: Single Nucleotide Polymorphism.

### Modeling TF targeting in melanoma

Melanoma progression and metastasis is known to be associated with many regulatory changes that alter patterns of gene expression [38], ultimately leading to phenotype switching to malignancy and drug resistance. These changes in expression can be tied to a variety of regulatory elements including transcription factor targeting, miRNA suppression of transcripts, and genomic and epigenetic changes. To demonstrate the utility of the netZoo framework for combining netZoo tools, we applied PANDA with LIONESS to model transcriptional regulation for individual samples in melanoma. This workflow allows us to understand regulatory changes in disease by estimating and analyzing sample-specific regulatory networks for the 76 melanoma cell lines available in CCLE and exploring a variety of disease-associated processes.

First, we used PANDA to generate an aggregate network across all CCLE cell lines, and we derived single-sample networks using LIONESS. Then, we used ANOVA to analyze the 76 melanoma networks to explore whether TF targeting scores, the sum of outgoing edge weights for each TF in the network [39], could be linked to methylation changes and copy number alterations. We defined hypermethylated and hypomethylated promoter sites as those having methylation status greater or less than three standard deviations from the mean (|z|>3) respectively; we considered a gene to be amplified if it had evidence of more than three copies in the genome and to be deleted if both copies are lost. We only computed the associations if they had at least three positive instances of the explanatory variable (for example, for a given gene at least three cell lines had a hypomethylation in that gene’s promoter) and corrected for multiple testing using a false discovery rate of less than 0.25 following the Benjamini-Hochberg procedure [40].

Among the top ten associations (Figure 2A), we found that the targeting by the TF melanocyte inducing transcription factor (MITF) was associated with changes in promoter methylation, in particular, we found a significant association between the MITF targeting score, computed as the TF weighted outdegree, and promoter hypermethylation of Discoidin, CUB and LCCL Domain Containing 2 (*DCBLD2*) (Figure 2A), a gene that has been suggested to trigger oncogenic processes in melanoma through Epidermal Growth Factor Receptor (*EGFR*) signaling [41]. This finding consistent with the identification of MITF as a key driver of melanoma [42, 43]. We also found that MITF targeting was associated with the deletion of Protein Tyrosine Phosphatase Non-Receptor Type 20 (*PTPN20*; Figure 2A), providing further evidence that disrupted signaling mediated by MITF regulation plays an important role in melanoma.

**Figure 2.**
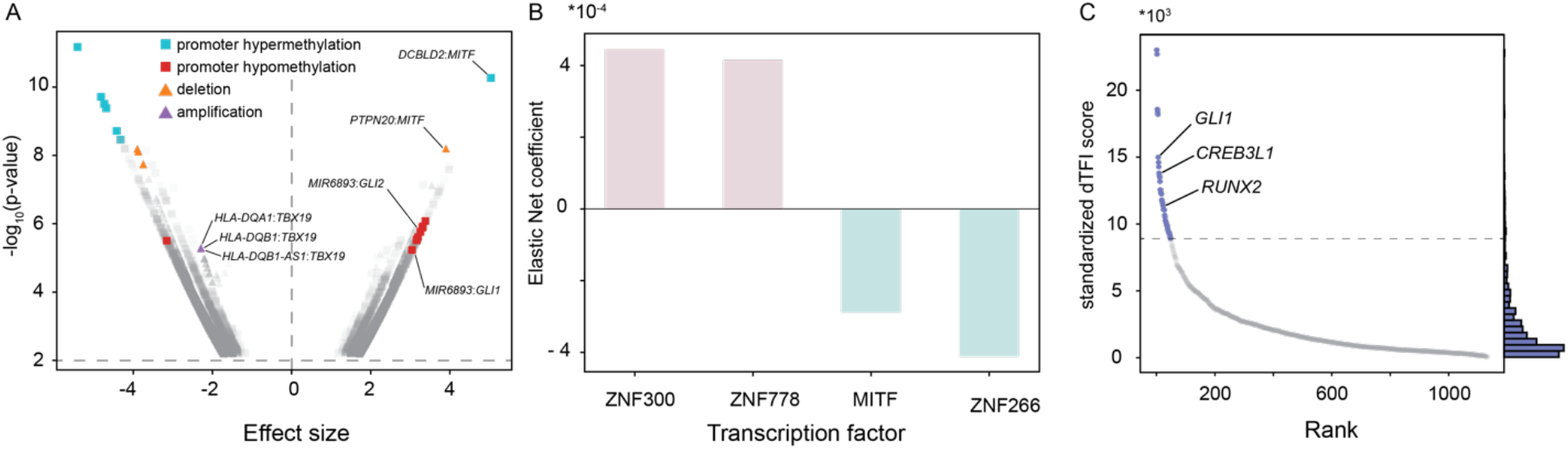
Modeling regulatory processes in melanoma using CCLE data. **A** Volcano plot of the ANOVA associations between TF targeting scores and promoter methylation and copy number statuses in melanoma cell lines. The 10 largest significant associations are colored in red and cyan for methylation status, and in orange and purple for copy number status. **B** Elastic net regression of Regorafenib cell viability on TF targeting scores in melanoma cell lines. The figure represents the two largest positive coefficients and two largest negative coefficients. **C** Differential TF involvement in the transition between primary melanoma cell line and a cell line derived from melanoma metastasis. The top 50 TFs are colored in blue.

The targeting by TFs glioma-associated oncogenes 1 and 2 (GLI1 and GLI2) was also significantly increased in melanoma; both TFs had previously been linked to drug resistance in melanoma cell lines [44]. In examining GLI1 and GLI2 targeting, we found it to be associated with promoter hypomethylation of *MIR6893*. According to TargetScan [10]), *MIR6893* regulates two related TFs, Glis Family Zinc Finger 1 and 2 (GLIS1 and GLIS2) and both have been reported to be involved in psoriasis [45], an inflammatory skin condition, which may indicate that they play a similar role in melanoma. Finally, examining additional significant associations (Figure 2A), we find a decrease of targeting by TBX19 to be associated with amplification of the *HLA-DBA1* and *HLA-DQB1* genes, both of which are known to be melanoma risk factors [46]. TBX19 itself has not been implicated in melanoma, but it has been linked to lymph node metastasis in colorectal cancer [47] and TBX2, another member of the T-Box family, is involved in melanoma proliferation [48].

We also tested whether TF targeting in the CCLE melanoma cell lines was associated with response to Regorafenib, a multi-kinase inhibitor that has been approved for treating metastatic colorectal cancer, advanced gastrointestinal stromal tumors, and advanced hepatocellular carcinoma. The drug has been shown to have a high affinity for BRAF [49], a kinase commonly mutated in metastatic melanoma, suggesting it may also show efficacy in treating melanoma. We performed Elastic Net regression [50] on TF targeting scores to test for cell viability following Regorafenib treatment [51] and among the largest variable importance, we found targeting by MITF to be negatively associated with cell viability (Figure 2B, Figure S1). This finding is consistent with studies that found that MITF loss to be associated with drug resistance [52] and underscores the multifunctional role that MITF appears to play in melanoma based on our analysis. However, other studies have implicated an increased activity of MITF in resistance to BRAF inhibitor treatment [53, 54]. Another TF, ZNF778, was also a strong predictor of Regorafenib sensitivity (Figure 2B, Figure S1); the ZNF778 promoter has also been found to be highly mutated in melanoma [55].

Finally, severe forms of melanoma are associated with a transition from a noninvasive to an invasive state [56] which can be driven by epithelial to mesenchymal transition (EMT). We used MONSTER to define a TF transition matrix that maps a nonmetastatic network for cell line GRN derived from a primary tumor (Depmap ID: ACH-000580) to one derived from a cell line derived from melanoma metastasis (Depmap ID: ACH-001569). We found that the TFs RUNX2, GLI1, and CREB3L1 were among those with the largest differential involvement score [24] (Figure 2C), indicating that they have the most profound changes in their regulatory targets as cells become metastatic. RUNX2 has been previously identified as a driver of epithelial to mesenchymal transition (EMT) processes and phenotype switching in melanoma [56]. CREB3LI has been reported to be activated in drug resistant cell lines [57] and GLI1 knockdown has been shown to increase sensitivity to Vemurafenib [44], an approved melanoma BRAF inhibitor. Collectively, the results from these analyses suggest a co-involvement of TFs associated with both drug resistance and cell state transition in invasive disease and highlight the promise of multi-kinase targeting [58].

### CCLE pan-cancer analysis reveals meaningful regulatory interactions

The CCLE cell lines are among the most widely studied model systems available in oncology research and include a large number of omic measurements as well as viability assays following drug challenges and gene knockdowns; we used DRAGON [16] to explore multi-omic associations captured in these data. DRAGON uses partial correlations with covariance shrinkage [59] to construct GGMs representing direct associations between measurements, accounting for the unique structure of each data type. We calculated DRAGON partial correlation networks between all pairwise sets of measurements on the CCLE cell lines, but we will focus on three sets of correlations in our discussion here: 1) miRNA levels and gene knockdown, 2) protein levels with metabolite levels, and 3) cell viability assays after drug exposure and gene knockdown screens.

In the first comparisons between miRNA expression and gene knockdowns, we assume that if a gene product is repressed by miRNA, then it is likely nonessential. We found that *MIR664* levels have a strong partial correlation with Glutathione-Disulfide Reductase (*GSR*) dependency, suggesting that *MIR664* post-transcriptionally regulates *GSR* (Figure 3A). This is consistent with annotation in the TargetScan database [10], which predicts *GSR* to be a target of *MIR664*, ranked 613/5387 with a context+ score of - 0.16. In our DRAGON analysis of metabolomic and proteomic data, we first found three glycolysis metabolites, Phosphoenolpyruvic acid, 3-phosphoglycerate, and glyceraldehyde 3P, were negatively partially correlated with Lactate Dehydrogenase-A (LDHA) protein levels (Figure S2, Figure 3B). This suggests that these metabolites are upstream of LDHA and indicates that glycolysis operates in the forward direction towards metabolite breakdown (Figure S2). Second, we found that fumarate/maleate levels, that are TCA cycle metabolites, were negatively partially correlated with LDHA (Figure 3B). These two observations suggest an activity of LDHA in the forward direction towards lactate production through aerobic fermentation (Figure S2). The activation of suboptimal pathways such as aerobic fermentation (the Warburg effect [60]) to produce energy is a hallmark of cancer and has been correlated with poor prognosis and drug resistance [61, 62]. This effect could be dominant in CCLE, since we did not filter for cell lines based on their metabolic phenotype.

**Figure 3.**
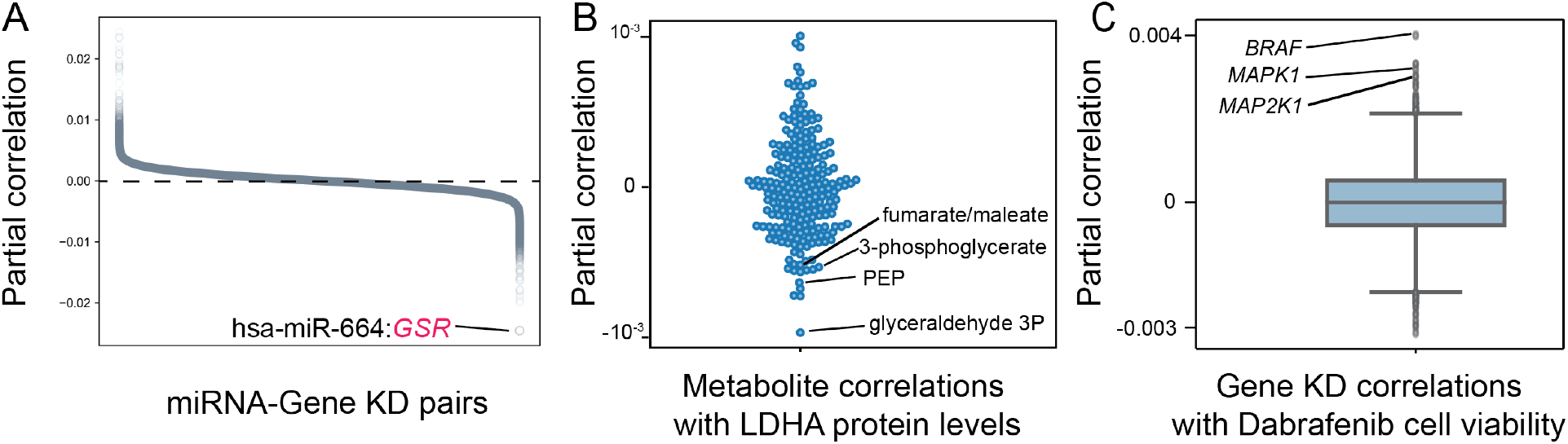
Pan-cancer analysis of regulatory interactions using DRAGON. **A** Partial correlation between miRNA levels and gene knockdown screen across all cancer cell lines. **B** Partial correlation of metabolite levels and LDHA protein levels. **C** Partial correlation between gene knockdown screens and Dabrafenib cell viability assays.

We also performed DRAGON analysis of cell viability assays after drug exposure and gene knockdown screens. Not surprisingly, we found that viability after exposure to Dabrafenib, a BRAF inhibitor, was highly correlated with *BRAF* knockdown. Dabrafenib cell viability was also correlated to *MAPK1* and *MAP2K1*, two genes that are downstream of BRAF in the MAPK signaling pathway. This finding is possibly due to compensatory mechanisms between functionally-related genes [63]. In the absence of these effects, the latter finding makes sense because although Dabrafenib is described as “selective” to BRAF [64], it has been shown to be active in cell lines with constitutively activated BRAF harboring the V600E activating mutation [65]; this subsequently triggers drug resistance by reactivating the MAPK pathway, particularly, MAPK1 and MAP2K1.

### An integrated CCLE multi-omic network portal

Having estimated DRAGON networks for additional pairwise combinations of measurements (Table S1) on the CCLE cell lines, we integrated these partial correlation networks from various omic types and created an online portal to allow exploration of the integrated relationships we discovered. Having built bipartite DRAGON networks between each pair of measurements, we systematically overlaid networks based on our understanding of the cascade of regulatory processes active in cells (Figure S3). First, promoter methylation status, copy number variation, histone marks, and miRNA partial correlations networks with gene expression were stacked to capture the multi-modal regulation of gene expression. Then, gene expression was linked to protein levels, which in turn was associated with cellular phenotypes represented by metabolite levels, drug sensitivity, and cell fitness resulting in a final genotype-to-phenotype map. To reduce the size of the network to the most relevant positive and negative associations, only the 2000 most positive correlations and the 2000 most negative correlations in each pairwise association in each of the bipartite networks were retained in the final multi-omic network.

The resulting integrated CCLE partial correlation network is available online (https://grand.networkmedicine.org/cclemap/) and can be queried to explore the biological associations contained within (Figure 4A). To illustrate the utility of this multi-tiered correlation network map we used it to examine the effect of copy number variation on gene expression. As expected, we found positive partial correlations between copy number and expression. For example, we not only found that *CDKN2A* and *CDKN2B* copy numbers have a positive partial correlation with *CDKN2A* and *CDKN2B* expression, respectively (Figure 4B), but that *CDKN2B* copy number is correlated with CDKN2A expression, which may reflect the fact that these two genes are adjacent in the genome. We also found negative partial correlations between copy number variation and gene expression. For example, *MIR378D1* copy number is negatively partially correlated with *TBC1D21* expression (Figure 4C), suggesting that *TBC1D21* may be repressed by *MIR378D1*. Although *TBC1D21* is not listed as a target of *MIR378D1* in miRDB, other members of the *TBC1* family, including *TBC1D12* (Target Score (TS) 66), *TBC1D16* (TS 61), and *TBC1D24* (TS 53), are among its predicted targets [66].

**Figure 4.**
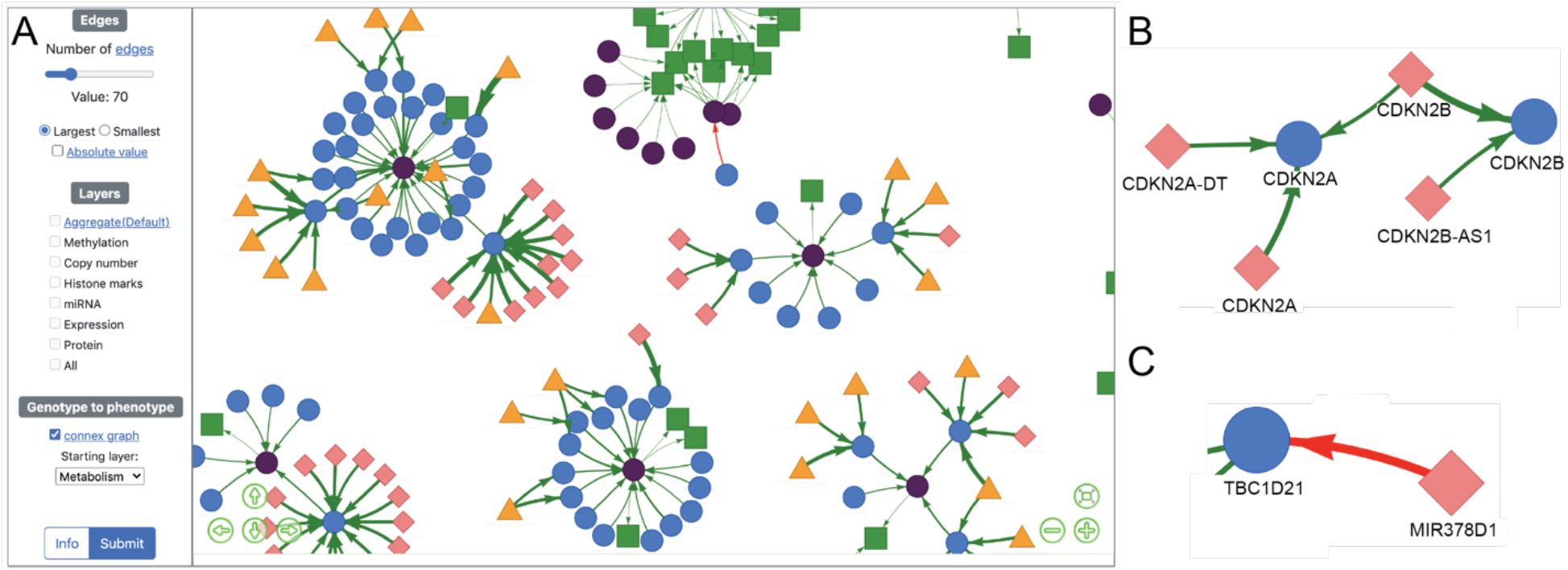
Multi-tiered CCLE map links genotype to cellular phenotypes. **A** Screenshot of the online resource accessible on https://grand.networkmedicine.org/cclemap/ that links promoter methylation (orange triangle), copy number variation (pink diamond), histone marks, miRNA levels, gene expression (blue circle), protein levels (purple circle), metabolite levels (green square), drug sensitivity, and cell fitness using DRAGON. **B** Positive partial correlations between copy number variation and gene expression of CDKN2A and CDKN2B. **C** Negative partial correlation between MIR378D1 levels and TBC1D21 expression.

### Creating a community ecosystem for collaborative software development

Development of netZoo has been driven through collaborative work involving users and developers at several academic institutions, all of whom are committed to open-source, community-driven tool development. A great deal of our work in harmonizing the code has been to facilitate reproducibility across implementation of related methods, to facilitate re-use of common methods for network inference, and to standardize input and output file formats to enable creation of network analysis pipelines.

The netZoo codebase is version-controlled in GitHub and implementations of most methods are available in R, Python, MATLAB, and C (Figure 5). Using a synchronized resource for code development avoids creating parallel branches and gives users access to tested and optimized tools that are up to date with the newest frameworks, particularly for the growing userbase in R and Python, as well as with third-party dependencies. The codebase includes additional helper functions for plotting and analysis, and GPU-accelerated implementations [67] for faster network inference across large numbers of samples. A continuous integration system called ZooKeeper runs unit tests using GitHub actions and a custom server to maintain the integrity of the software and update dependencies to third-party software. In addition, contributions from the community follow a fork-branch model and are tested through ZooKeeper before being added to the core codebase. Finally, users can access a set of cloud-based Jupyter notebook use cases and tutorials hosted through Netbooks [28] and visualize a set of networks hosted in the GRAND database [29].

**Figure 5.**
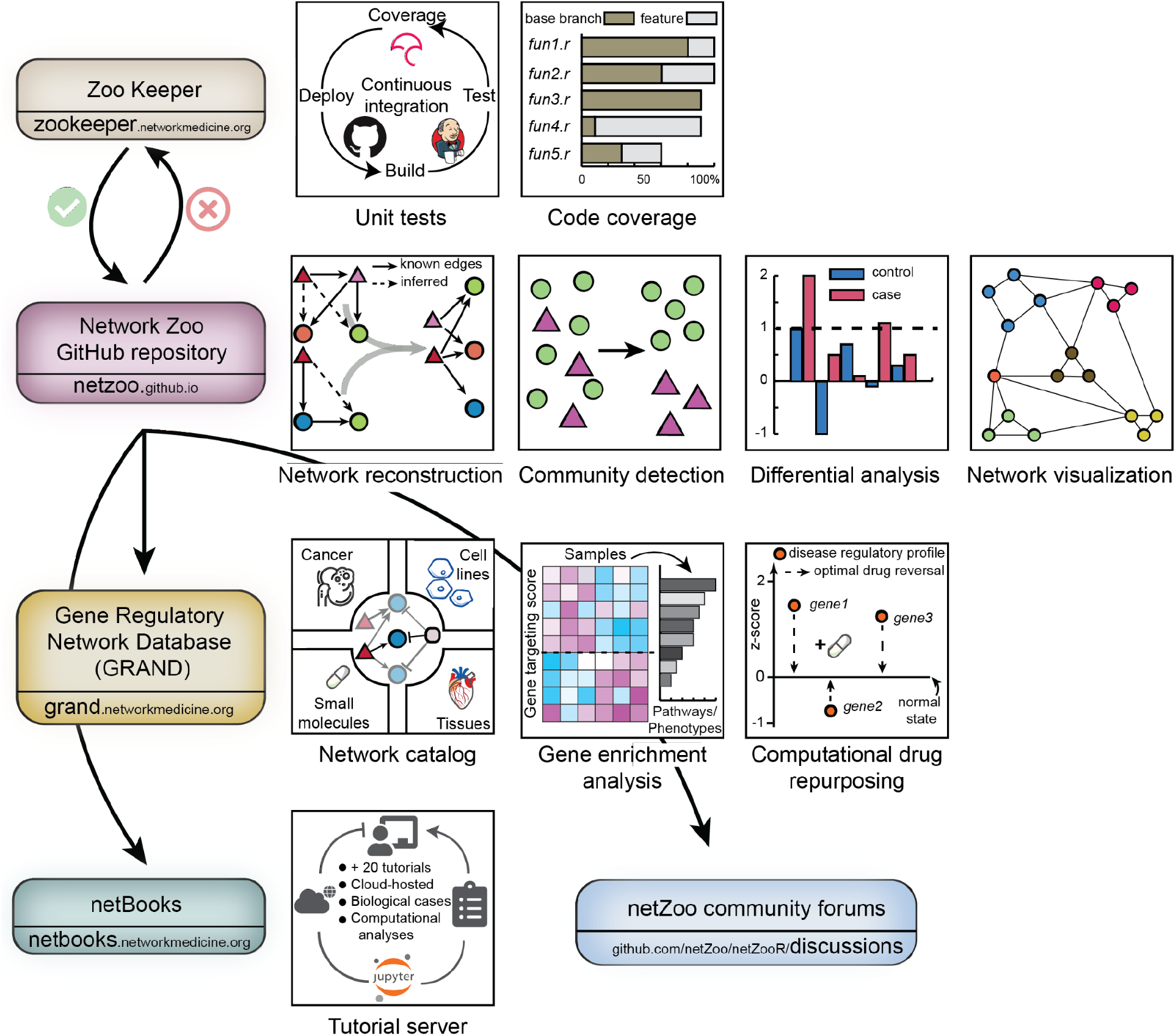
netZoo ecosystem. The codebase is hosted on GitHub and is regularly tested through a continuous integration system called ZooKeeper. Networks generated by netZoo tools are hosted in the GRAND database. Cloud-hosted use cases and tutorials are available through a JupyterHub server called Netbooks. GitHub discussions and issues provide a forum for discussion and exchange within the community.

## Conclusions

We developed netZoo as an open-source platform for the inference and analysis of biological networks, including multi-omic gene regulatory and partial correlation networks. We accomplished this by standardizing the implementations of software tools built on a common conceptual framework, in line with recent similar efforts [68, 69], and building an ecosystem for sharing of use cases, hosting networks, and continued development and maintenance which is essential for software accuracy [70]. We will continue to expand netZoo (Figure S4), adding new methods [71–73] and improving implementations of the existing tools, as well as building interfaces to allow methods to be combined appropriately. We welcome community participation in methods development and are committed to the broad use of the tools available within netZoo.

## Methods

### netZoo applications using the Cancer Cell Line Encyclopedia

The CCLE project characterized more than a thousand cell lines from 35 cancer types, measuring gene and miRNA expression, promoter methylation status, copy number variation, protein and metabolite levels. Phenotypic data are available from the PRISM project on viability of these cell lines following drug exposure [74] and from cell fitness screens available through the dependency map [75]. For the 1,376 CCLE cell lines that had transcriptomic measurements, we inferred GRNs using PANDA and LIONESS algorithms available in netZooPy v0.8.1 and used these for various analyses.

As input to PANDA network inference process, we began with a TF-to-gene prior regulatory network computed by running FIMO [33] scans of 1,149 TF motifs from CIS-BP (v1.94d; [5]) in the promoter region of genes (defined as 1kb downstream of each gene’s transcription start site) in the reference human genome sequence (hg38); we adjusted the TF-gene pair by combining two previously-suggested scores [76, 77]. The modified score (*s*) integrates the distance between the detected motif and the TSS with the significance of motif assignment as follows:

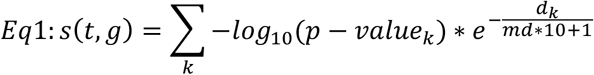

where *t* is a transcription factor, *g* is a target gene, *k* is the number of binding sites of *t* identified in the promoter region of *g, d_k_* denotes the distance of *t*’s biding site *k* to TSS of *g, md* the median of all the distances *d*, and *p-valuek* the significance of assignment of binding site *k*. We used as inputs a TF PPI network derived from the STRING database [2] (using the aggregate score for human interactions only and scaling them between 0 and 1) and a gene-gene correlation network based on gene expression data, as preprocessed in CCLE. The resulting PANDA network includes regulatory associations between 1,132 TFs and 18,560 genes using the ‘mode’ parameter to intersection which take the intersection TF and genes between the three input networks. Then, we used LIONESS to infer regulatory networks for each of the 1,376 cell lines; all networks can be found in the GRAND database (https://grand.networkmedicine.org/cell/). We also computed TF targeting scores [78] by computing the weighted outdegree for each TF in each cell line specific network.

### Modeling TF targeting associations in melanoma

To find associations between TF targeting and promoter methylation status and copy number variation status, we selected 76 melanoma CCLE cell lines and we computed the significance of associations using ANOVA. We considered a gene to be amplified if it had more than three copies and to be deleted if both copies are lost. Promoters were defined in CCLE as the 1kb region downstream of the gene’s transcriptional start site (TSS). Promoter hypermethylation was computed by z-scoring the methylation matrix across all samples and taking promoters that had a z-score larger than three. Similarly, a promoter was considered hypomethylated if it had a z-score less than three.

In all melanoma cell lines, for each modality (promoter hypomethylation, promoter hypermethylation, gene amplification, and gene deletion) and for each gene, we built an ANOVA model using TF targeting as the response variable across all melanoma cell lines while the status of that gene (either promoter methylation or copy number status) was the explanatory variable (with positive instances for methylated promoters/amplified genes and negative instances for nonmethylated promoters/nonamplified genes) along with an additional factor correcting for the cell lineage. We only computed the associations if they had at least three positive instances of the explanatory variable. We corrected for multiple testing by taking associations that had less than 25% false discovery rate.

To predict drug response using TF targeting, we conducted a linear regression with Elastic Net [50] regularization using an equal weight of 0.5 for L1 and L2 penalties using Regorafenib cell viability assays in melanoma cell lines as a response variable and the targeting scores of 1,132 TFs (Table S2) as the explanatory variable.

Finally, to model EMT in melanoma, we used MONSTER on two LIONESS networks of melanoma cancer cell lines, one representing a primary tumor (Depmap ID: ACH-000580) as the initial state and the other a metastasis cell line (Depmap ID: ACH-001569) as the end state. We modified the original implementation of MONSTER that implements its own network reconstruction procedure to take any input network, such as LIONESS networks in our case. MONSTER identifies differentially-involved TFs in the transition by shuffling the columns of the initial and final state adjacency matrices 1000 times to build a null distribution, which is then used to compute a standardized differential TF involvement score by scaling the obtained scores by those of the null distribution.

### Computing CCLE multi-omic associations

We used DRAGON to compute partial correlations between multi-omic data of CCLE cell lines. In particular, we computed partial correlations between the three following data-type pairs across all CCLE cell lines: 1) miRNA levels and gene knockdown, 2) protein levels and metabolite levels, and 3) cell viability assays after drug exposure and gene knockdown screens. DRAGON builds a GGM that implements covariance shrinkage with omic-specific tuning parameters, a novel addition to covariance shrinkage that enables DRAGON to account for varying data structures and sparsities of different multi-omic layers [59]. All variables were standardized to have a mean of 0 and a standard deviation of 1 before calling DRAGON.

### CCLE pan-cancer map

To enable further exploration and discovery of biological associations, we built an online resource representing a multi-tiered regulatory network. First, to build a pan-cancer multi-tiered network that connects the genotype to cellular phenotypes, we extended DRAGON networks from modeling pairwise interactions between two omics to a multi-omic network by sequentially adding a new layer to an initial pairwise DRAGON network. In addition, since DRAGON networks are undirected, we added direction based on our understanding of how biological elements interact with each other. For example, gene expression nodes are upstream of protein levels nodes and metabolite nodes. To facilitate browsing and limit exploration to potentially causal associations that best reflect our understanding of how different data types link to one another in cellular biology, our approach was to prune edges between the same node type to build bipartite DRAGON networks between each pair of genomic modalities. In particular, promoter methylation status, copy number variation, histone marks, and miRNA were linked to gene expression in a pairwise fashion. Then, gene expression was linked to protein levels, which in turn was associated with cellular phenotypes represented by metabolite levels, drug sensitivity, and cell fitness. Only the 2000 most positive correlations and the 2000 most negative correlations in each pairwise association were considered in the final network. The CCLE online pan-cancer map was built using Vis.js (v8.5.2) and can be queried for biological associations using user input queries at https://grand.networkmedicine.org/cclemap.

For all analysis presented in this work, we used the following releases of CCLE data: Promoter methylation data of 2018/10/22, histone marks data of 2018/11/30, miRNA expression data of 2018/11/03, metabolite levels data [32] of 2019/05/02. Cell viability assays were taken from the 19Q4 release of PRISM [74]. Cell fitness screens were taken from the 21Q1 release of project Achilles. Gene expression and copy number variation were taken from the 21Q1 release of the Dependency Map. Protein levels [31] were taken from the 2020/01 version of CCLE.

### Software package

All analyses were performed using netZooPy v0.8.1, the Python distribution of the netZoo (netzoo.github.io). NetZoo tools are implemented in R, Python, MATLAB, and C. netZooR v1.0 is currently implemented in R v4.1 and available through GitHub (https://github.com/netZoo/netZooR) and Bioconductor (https://bioconductor.org/packages/netZooR) and includes PANDA, LIONESS, CONDOR, ALPACA, SAMBAR, MONSTER, OTTER, CRANE, EGRET, and YARN. netZooPy v0.8.1 is implemented in Python v3.9 and includes PANDA, LIONESS, PUMA, CONDOR, SAMBAR, OTTER, and DRAGON. netZooM v0.5.2 is implemented in MATLAB 2020b (The Mathworks, Natick, MA, USA) and includes PANDA, LIONESS, PUMA, SPIDER, and OTTER. netZooC v0.2 implements PANDA and PUMA.

## Supplementary Information

Additional file 1 contains four supplementary figures and two supplementary tables.

## Acknowledgements

The authors would like to acknowledge Yunhao Huo for assistance with graphical design.

## Authors’ contributions

MBG, TW, CMLR, VF, and JQ designed the project and maintained the software distribution. KG, MLK, AS, JP, DW, MP, JL, RB, MA, DS, and JNP developed the methods. KG, GC, DVI, AM, RB, DS, JNP implemented methods in a different programming language. AM, DM, CYC, DD, and KS implemented new features. AS, MLK, GC, JP, DW, DM, MP, MA, DS, JNP wrote vignettes and tutorials. MBG and JQ wrote the manuscript with input from all the authors. All authors read and approved the final manuscript.

## Authors’ information

Twitter handles: Marouen Ben Guebila (@marouenbg), Camila M. Lopes-Ramos (@camilamlopes), Viola Fanfani (@violafanfani), Daniel Schlauch (@dschlauch), Joseph N. Paulson (@josephnpaulson), Abhijeet Sonawane (@abhijeetrs), Genis Calderer (@genisott), David van Ijzendoorn (@vanijzen), Daniel Morgan (@dcolinmorgan), Alex Song (@sqsq3178), Megha Padi (@megha_padi), Marieke L. Kuijjer (@mkuijjer), John Quackenbush (@johnquackenbush)

## Availability of data and materials

netZoo tools are available at http://netzoo.github.io. CCLE cell line GRNs can be downloaded at https://grand.networkmedicine.org/cell/ and the CCLE multi-tiered map can be accessed at https://grand.networkmedicine.org/cclemap/.

## Declarations

### Ethics approval and consent to participate

Not applicable

### Consent for publication

Not applicable

