## Additional file 1 for "The Network Zoo: a multilingual package for the inference and analysis of biological networks"

^2^Present address: Biology Department, Boston College, Chestnut Hill, MA, USA

^4^Present address: Lineberger Comprehensive Cancer Center, University of North Carolina at Chapel Hill, NC, USA

^5^Present address: CISPA Helmholtz Center for Information Security, Saarbrücken, Germany

^6^Present address: Genospace, LLC, Boston, MA, USA

^8^Present address: Department of Medical Bioinformatics, University Medical Center Göttingen, Göttingen, Germany

^9^Present address: Center for Interdisciplinary Cardiovascular Sciences, Division of Cardiovascular Medicine, Department of Medicine, Brigham and Women’s Hospital, Boston, MA, USA

^11^Present address: Monoceros Biosystems, LLC, San Diego, CA, USA

^14^Present address: Department of Pathology, Stanford University School of Medicine, CA, USA

^15^Present address: Hong Kong University, School of Biomedical Sciences, Honk Kong

^18^Present address: Institute of Biomedical Informatics, National Yang Ming Chiao Tung University, Taipei 112, Taiwan

^19^Present address: Computational Biology Department, Carnegie Mellon University, Pittsburgh, PA, USA

**Supplementary figures**

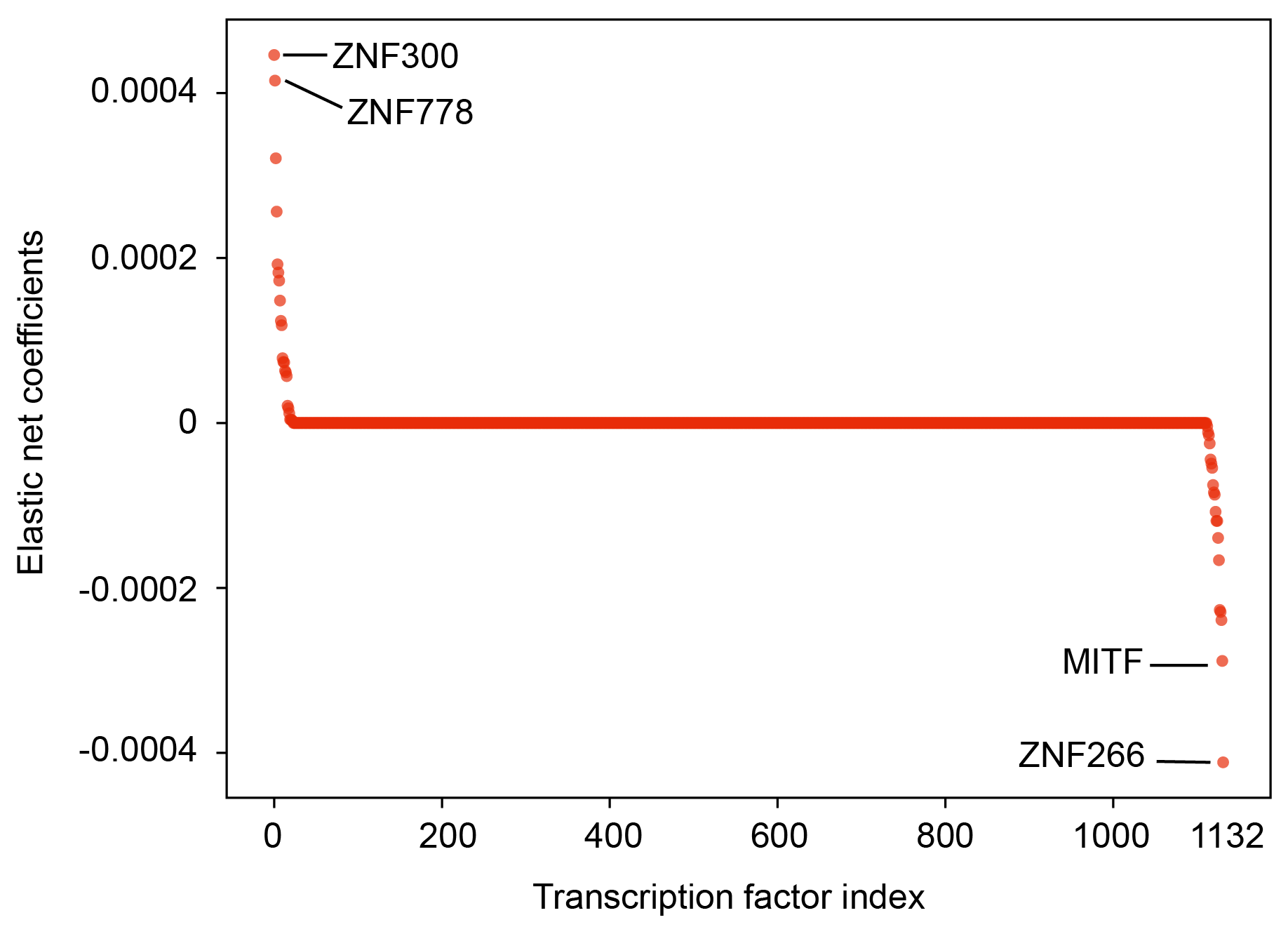

**Figure S1** Elastic Net coefficients of Regorafenib drug sensitivity regression on TF targeting. The analysis includes all 1,132 TFs modeled in the GRNs of 76 melanoma cell lines. The tails of this distribution are represented in Figure 2B.

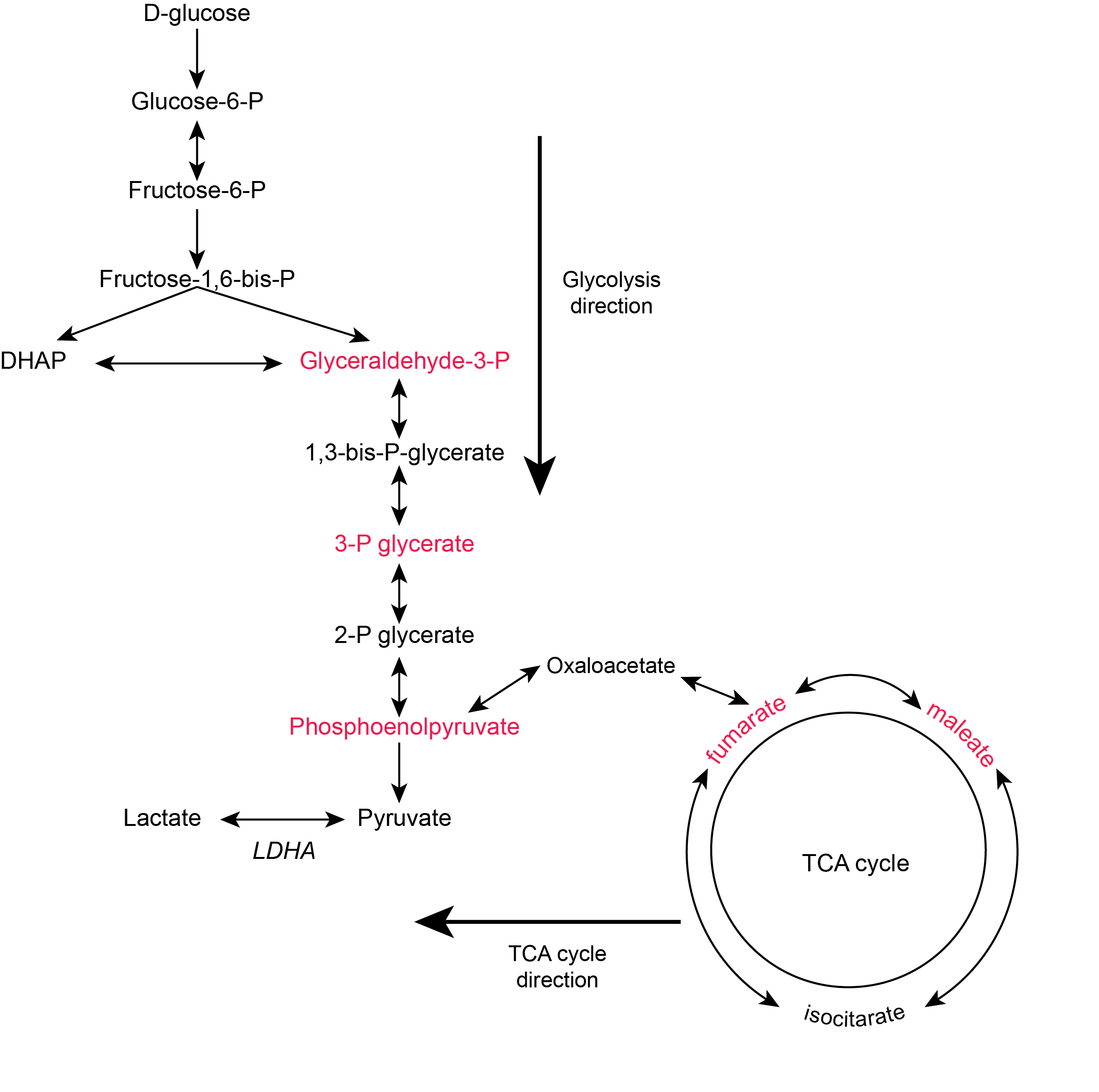

**Figure S2** Predicted glycolysis and TCA cycle directions based on metabolite and enzyme levels. Red-colored metabolites are negatively correlated with *LDHA* enzyme levels in CCLE cell lines which suggests that Glycolysis operates in the forward direction and that TCA cycle does not break down the product of glycolysis, thereby a switch to lactate production pathway.

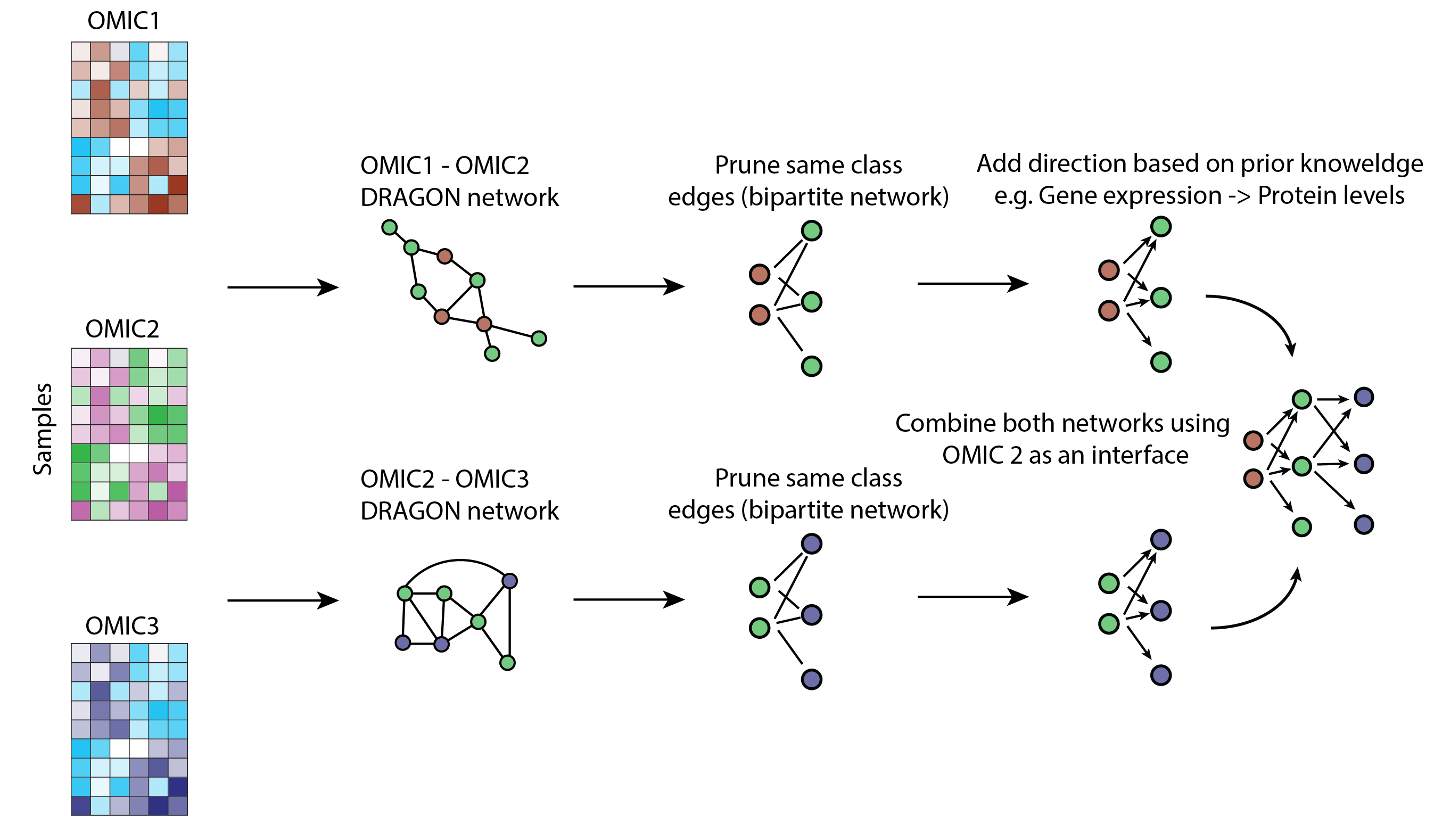

**Figure S3** Reconstruction of a multi-omic partial correlation network using DRAGON. The example illustrates the creation of a tripartite network. DRAGON generates unipartite undirected network between two pairs of “omics”. To reflect prior biological knowledge, we prune same class edges and add edges directions. Then, we combine two networks by using the intersecting ‘omic’ as an interface.

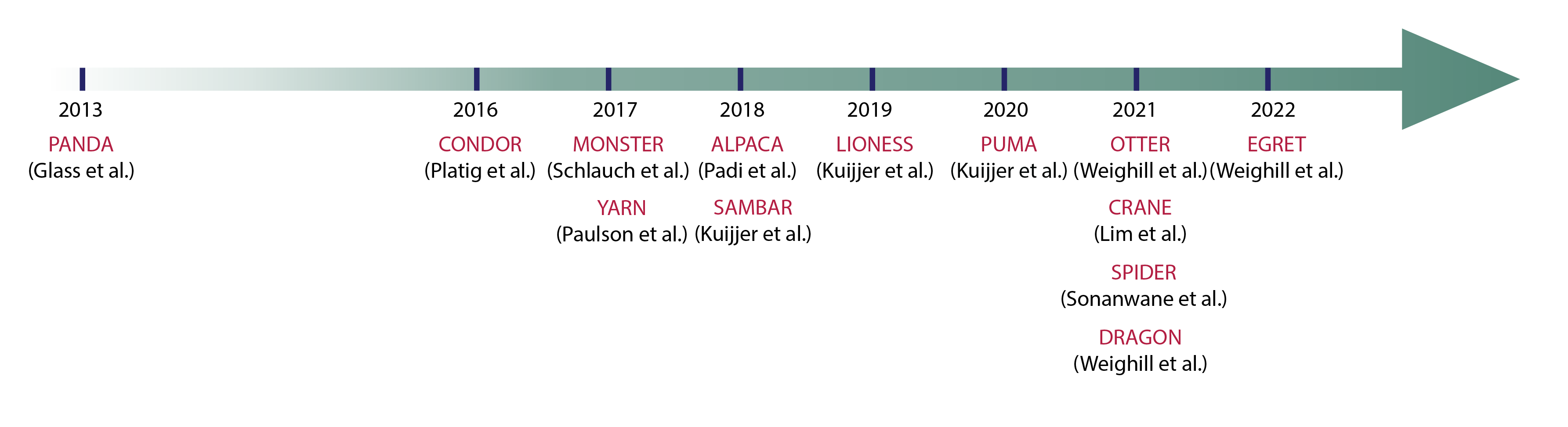
**Figure S4** Timeline of netZoo methods’ publications.

**Supplementary tables**

**Table S1** – Pairwise combinations of multi-omic data to build a CCLE integrated partial correlation network

| Pair | Omic 1 | Omic 2 |
| --- | --- | --- |
| 1 | Promoter methylation | Gene expression |
| 2 | Copy number variation | Gene expression |
| 3 | Histone marks | Gene expression |
| 4 | miRNA expression | Gene expression |
| 5 | Gene expression | Protein levels |
| 6 | Protein levels | Metabolite levels |
| 7 | Protein levels | Drug sensitivity (Cell viability) |
| 8 | Protein levels | Cell fitness (CRISPR KO) |

**Table S2** – Genes names of 1,132 TFs modeled in CCLE GRNs.

| **ID** | **TF** |
| --- | --- |
| 1 | AHR |
| 2 | AHRR |
| 3 | AIRE |
| 4 | ALX1 |
| 5 | ALX3 |
| 6 | ALX4 |
| 7 | ANHX |
| 8 | AR |
| 9 | ARGFX |
| 10 | ARID3A |
| 11 | ARID3B |
| 12 | ARID3C |
| 13 | ARID5A |
| 14 | ARID5B |
| 15 | ARNT |
| 16 | ARNT2 |
| 17 | ARNTL |
| 18 | ARNTL2 |
| 19 | ARX |
| 20 | ASCL1 |
| 21 | ASCL2 |
| 22 | ASCL3 |
| 23 | ASCL4 |
| 24 | ASCL5 |
| 25 | ATF1 |
| 26 | ATF2 |
| 27 | ATF3 |
| 28 | ATF4 |
| 29 | ATF5 |
| 30 | ATF6 |
| 31 | ATF6B |
| 32 | ATF7 |
| 33 | ATOH1 |
| 34 | ATOH7 |
| 35 | ATOH8 |
| 36 | BACH1 |
| 37 | BACH2 |
| 38 | BARHL1 |
| 39 | BARHL2 |
| 40 | BARX1 |
| 41 | BARX2 |
| 42 | BATF |
| 43 | BATF3 |
| 44 | BBX |
| 45 | BCL11A |
| 46 | BCL11B |
| 47 | BCL6 |
| 48 | BCL6B |
| 49 | BHLHA15 |
| 50 | BHLHE22 |
| 51 | BHLHE23 |
| 52 | BHLHE40 |
| 53 | BHLHE41 |
| 54 | BPTF |
| 55 | BSX |
| 56 | CDC5L |
| 57 | CDX1 |
| 58 | CDX2 |
| 59 | CDX4 |
| 60 | CEBPA |
| 61 | CEBPB |
| 62 | CEBPD |
| 63 | CEBPE |
| 64 | CEBPG |
| 65 | CEBPZ |
| 66 | CENPB |
| 67 | CIC |
| 68 | CLOCK |
| 69 | CPEB1 |
| 70 | CREB1 |
| 71 | CREB3 |
| 72 | CREB3L1 |
| 73 | CREB3L2 |
| 74 | CREB3L4 |
| 75 | CREB5 |
| 76 | CREBL2 |
| 77 | CREBZF |
| 78 | CREM |
| 79 | CRX |
| 80 | CTCF |
| 81 | CTCFL |
| 82 | CUX1 |
| 83 | CUX2 |
| 84 | CXXC1 |
| 85 | DBP |
| 86 | DBX1 |
| 87 | DBX2 |
| 88 | DDIT3 |
| 89 | DLX1 |
| 90 | DLX2 |
| 91 | DLX3 |
| 92 | DLX4 |
| 93 | DLX5 |
| 94 | DLX6 |
| 95 | DMBX1 |
| 96 | DMRT1 |
| 97 | DMRT2 |
| 98 | DMRT3 |
| 99 | DMRTA1 |
| 100 | DMRTA2 |
| 101 | DMRTC2 |
| 102 | DNMT1 |
| 103 | DPF1 |
| 104 | DPF3 |
| 105 | DPRX |
| 106 | DRGX |
| 107 | DUX4 |
| 108 | DUXA |
| 109 | E2F1 |
| 110 | E2F2 |
| 111 | E2F3 |
| 112 | E2F4 |
| 113 | E2F5 |
| 114 | E2F6 |
| 115 | E2F7 |
| 116 | E2F8 |
| 117 | E4F1 |
| 118 | EBF1 |
| 119 | EBF2 |
| 120 | EBF3 |
| 121 | EBF4 |
| 122 | EGR1 |
| 123 | EGR2 |
| 124 | EGR3 |
| 125 | EGR4 |
| 126 | EHF |
| 127 | ELF1 |
| 128 | ELF2 |
| 129 | ELF3 |
| 130 | ELF4 |
| 131 | ELF5 |
| 132 | ELK1 |
| 133 | ELK3 |
| 134 | ELK4 |
| 135 | EMX1 |
| 136 | EMX2 |
| 137 | EN1 |
| 138 | EN2 |
| 139 | EOMES |
| 140 | EPAS1 |
| 141 | ERF |
| 142 | ERG |
| 143 | ESR1 |
| 144 | ESR2 |
| 145 | ESRRA |
| 146 | ESRRB |
| 147 | ESRRG |
| 148 | ESX1 |
| 149 | ETS1 |
| 150 | ETS2 |
| 151 | ETV1 |
| 152 | ETV2 |
| 153 | ETV3 |
| 154 | ETV3L |
| 155 | ETV4 |
| 156 | ETV5 |
| 157 | ETV6 |
| 158 | ETV7 |
| 159 | EVX1 |
| 160 | EVX2 |
| 161 | FERD3L |
| 162 | FEV |
| 163 | FEZF1 |
| 164 | FEZF2 |
| 165 | FIGLA |
| 166 | FLI1 |
| 167 | FOS |
| 168 | FOSB |
| 169 | FOSL1 |
| 170 | FOSL2 |
| 171 | FOXA1 |
| 172 | FOXA2 |
| 173 | FOXA3 |
| 174 | FOXB1 |
| 175 | FOXB2 |
| 176 | FOXC1 |
| 177 | FOXC2 |
| 178 | FOXD1 |
| 179 | FOXD2 |
| 180 | FOXD3 |
| 181 | FOXD4 |
| 182 | FOXD4L1 |
| 183 | FOXD4L3 |
| 184 | FOXD4L4 |
| 185 | FOXD4L5 |
| 186 | FOXD4L6 |
| 187 | FOXE1 |
| 188 | FOXE3 |
| 189 | FOXF1 |
| 190 | FOXF2 |
| 191 | FOXG1 |
| 192 | FOXH1 |
| 193 | FOXI1 |
| 194 | FOXI2 |
| 195 | FOXI3 |
| 196 | FOXJ1 |
| 197 | FOXJ2 |
| 198 | FOXJ3 |
| 199 | FOXK1 |
| 200 | FOXK2 |
| 201 | FOXL1 |
| 202 | FOXL2 |
| 203 | FOXM1 |
| 204 | FOXN1 |
| 205 | FOXN2 |
| 206 | FOXN3 |
| 207 | FOXN4 |
| 208 | FOXO1 |
| 209 | FOXO3 |
| 210 | FOXO4 |
| 211 | FOXO6 |
| 212 | FOXP1 |
| 213 | FOXP2 |
| 214 | FOXP3 |
| 215 | FOXP4 |
| 216 | FOXQ1 |
| 217 | FOXR1 |
| 218 | FOXR2 |
| 219 | FOXS1 |
| 220 | GABPA |
| 221 | GATA1 |
| 222 | GATA2 |
| 223 | GATA3 |
| 224 | GATA4 |
| 225 | GATA5 |
| 226 | GATA6 |
| 227 | GBX1 |
| 228 | GBX2 |
| 229 | GCM1 |
| 230 | GCM2 |
| 231 | GFI1 |
| 232 | GFI1B |
| 233 | GLI1 |
| 234 | GLI2 |
| 235 | GLI3 |
| 236 | GLI4 |
| 237 | GLIS1 |
| 238 | GLIS2 |
| 239 | GLIS3 |
| 240 | GMEB1 |
| 241 | GMEB2 |
| 242 | GRHL1 |
| 243 | GRHL2 |
| 244 | GSC |
| 245 | GSC2 |
| 246 | GSX1 |
| 247 | GSX2 |
| 248 | GTF3A |
| 249 | HAND1 |
| 250 | HAND2 |
| 251 | HBP1 |
| 252 | HDX |
| 253 | HELT |
| 254 | HES1 |
| 255 | HES2 |
| 256 | HES3 |
| 257 | HES4 |
| 258 | HES5 |
| 259 | HES6 |
| 260 | HES7 |
| 261 | HESX1 |
| 262 | HEY1 |
| 263 | HEY2 |
| 264 | HEYL |
| 265 | HHEX |
| 266 | HIC1 |
| 267 | HIC2 |
| 268 | HIF1A |
| 269 | HIF3A |
| 270 | HINFP |
| 271 | HIVEP1 |
| 272 | HIVEP2 |
| 273 | HIVEP3 |
| 274 | HKR1 |
| 275 | HLF |
| 276 | HLX |
| 277 | HMBOX1 |
| 278 | HMG20B |
| 279 | HMGA1 |
| 280 | HMGA2 |
| 281 | HMX1 |
| 282 | HMX2 |
| 283 | HMX3 |
| 284 | HNF1A |
| 285 | HNF1B |
| 286 | HNF4A |
| 287 | HNF4G |
| 288 | HOMEZ |
| 289 | HOXA1 |
| 290 | HOXA10 |
| 291 | HOXA11 |
| 292 | HOXA13 |
| 293 | HOXA2 |
| 294 | HOXA3 |
| 295 | HOXA4 |
| 296 | HOXA5 |
| 297 | HOXA6 |
| 298 | HOXA7 |
| 299 | HOXA9 |
| 300 | HOXB1 |
| 301 | HOXB13 |
| 302 | HOXB2 |
| 303 | HOXB3 |
| 304 | HOXB4 |
| 305 | HOXB5 |
| 306 | HOXB6 |
| 307 | HOXB7 |
| 308 | HOXB8 |
| 309 | HOXB9 |
| 310 | HOXC10 |
| 311 | HOXC11 |
| 312 | HOXC12 |
| 313 | HOXC13 |
| 314 | HOXC4 |
| 315 | HOXC5 |
| 316 | HOXC6 |
| 317 | HOXC8 |
| 318 | HOXC9 |
| 319 | HOXD1 |
| 320 | HOXD10 |
| 321 | HOXD11 |
| 322 | HOXD12 |
| 323 | HOXD13 |
| 324 | HOXD3 |
| 325 | HOXD4 |
| 326 | HOXD8 |
| 327 | HOXD9 |
| 328 | HSF1 |
| 329 | HSF2 |
| 330 | HSF4 |
| 331 | HSF5 |
| 332 | IKZF1 |
| 333 | IKZF2 |
| 334 | IKZF3 |
| 335 | IKZF4 |
| 336 | INSM1 |
| 337 | IRF1 |
| 338 | IRF2 |
| 339 | IRF3 |
| 340 | IRF4 |
| 341 | IRF5 |
| 342 | IRF6 |
| 343 | IRF7 |
| 344 | IRF8 |
| 345 | IRF9 |
| 346 | IRX1 |
| 347 | IRX2 |
| 348 | IRX3 |
| 349 | IRX4 |
| 350 | IRX5 |
| 351 | IRX6 |
| 352 | ISL1 |
| 353 | ISL2 |
| 354 | ISX |
| 355 | JDP2 |
| 356 | JUN |
| 357 | JUNB |
| 358 | JUND |
| 359 | KDM2B |
| 360 | KLF1 |
| 361 | KLF10 |
| 362 | KLF11 |
| 363 | KLF12 |
| 364 | KLF13 |
| 365 | KLF14 |
| 366 | KLF15 |
| 367 | KLF16 |
| 368 | KLF17 |
| 369 | KLF2 |
| 370 | KLF3 |
| 371 | KLF4 |
| 372 | KLF5 |
| 373 | KLF6 |
| 374 | KLF7 |
| 375 | KLF8 |
| 376 | KLF9 |
| 377 | KMT2A |
| 378 | LBX1 |
| 379 | LBX2 |
| 380 | LCOR |
| 381 | LEF1 |
| 382 | LHX1 |
| 383 | LHX2 |
| 384 | LHX3 |
| 385 | LHX4 |
| 386 | LHX5 |
| 387 | LHX6 |
| 388 | LHX8 |
| 389 | LHX9 |
| 390 | LIN28A |
| 391 | LIN28B |
| 392 | LIN54 |
| 393 | LMX1A |
| 394 | LMX1B |
| 395 | LYL1 |
| 396 | MAF |
| 397 | MAFA |
| 398 | MAFB |
| 399 | MAFF |
| 400 | MAFG |
| 401 | MAFK |
| 402 | MAX |
| 403 | MAZ |
| 404 | MBD2 |
| 405 | MBNL2 |
| 406 | MECOM |
| 407 | MECP2 |
| 408 | MEF2A |
| 409 | MEF2B |
| 410 | MEF2C |
| 411 | MEF2D |
| 412 | MEIS1 |
| 413 | MEIS2 |
| 414 | MEIS3 |
| 415 | MEOX1 |
| 416 | MEOX2 |
| 417 | MESP1 |
| 418 | MESP2 |
| 419 | MGA |
| 420 | MITF |
| 421 | MIXL1 |
| 422 | MLX |
| 423 | MLXIP |
| 424 | MLXIPL |
| 425 | MNT |
| 426 | MNX1 |
| 427 | MSC |
| 428 | MSGN1 |
| 429 | MSX1 |
| 430 | MSX2 |
| 431 | MTF1 |
| 432 | MXD1 |
| 433 | MXD3 |
| 434 | MXD4 |
| 435 | MXI1 |
| 436 | MYB |
| 437 | MYBL1 |
| 438 | MYBL2 |
| 439 | MYC |
| 440 | MYCL |
| 441 | MYCN |
| 442 | MYF5 |
| 443 | MYF6 |
| 444 | MYNN |
| 445 | MYOD1 |
| 446 | MYOG |
| 447 | MYRF |
| 448 | MYT1L |
| 449 | MZF1 |
| 450 | NAIF1 |
| 451 | NANOG |
| 452 | NEUROD1 |
| 453 | NEUROD2 |
| 454 | NEUROD4 |
| 455 | NEUROD6 |
| 456 | NEUROG1 |
| 457 | NEUROG2 |
| 458 | NEUROG3 |
| 459 | NFAT5 |
| 460 | NFATC1 |
| 461 | NFATC2 |
| 462 | NFATC3 |
| 463 | NFATC4 |
| 464 | NFE2 |
| 465 | NFE2L1 |
| 466 | NFE2L2 |
| 467 | NFE2L3 |
| 468 | NFIA |
| 469 | NFIB |
| 470 | NFIC |
| 471 | NFIL3 |
| 472 | NFIX |
| 473 | NFKB1 |
| 474 | NFKB2 |
| 475 | NFYA |
| 476 | NFYB |
| 477 | NFYC |
| 478 | NHLH1 |
| 479 | NHLH2 |
| 480 | NKX1-1 |
| 481 | NKX1-2 |
| 482 | NKX2-1 |
| 483 | NKX2-2 |
| 484 | NKX2-3 |
| 485 | NKX2-4 |
| 486 | NKX2-5 |
| 487 | NKX2-6 |
| 488 | NKX2-8 |
| 489 | NKX3-1 |
| 490 | NKX3-2 |
| 491 | NKX6-1 |
| 492 | NKX6-2 |
| 493 | NKX6-3 |
| 494 | NOBOX |
| 495 | NOTO |
| 496 | NPAS1 |
| 497 | NPAS2 |
| 498 | NPAS3 |
| 499 | NR0B1 |
| 500 | NR1D1 |
| 501 | NR1D2 |
| 502 | NR1H2 |
| 503 | NR1H3 |
| 504 | NR1H4 |
| 505 | NR1I2 |
| 506 | NR1I3 |
| 507 | NR2C1 |
| 508 | NR2C2 |
| 509 | NR2E1 |
| 510 | NR2E3 |
| 511 | NR2F1 |
| 512 | NR2F2 |
| 513 | NR2F6 |
| 514 | NR3C1 |
| 515 | NR3C2 |
| 516 | NR4A1 |
| 517 | NR4A2 |
| 518 | NR4A3 |
| 519 | NR5A1 |
| 520 | NR5A2 |
| 521 | NR6A1 |
| 522 | NRF1 |
| 523 | NRL |
| 524 | OLIG1 |
| 525 | OLIG2 |
| 526 | OLIG3 |
| 527 | ONECUT1 |
| 528 | ONECUT2 |
| 529 | ONECUT3 |
| 530 | OSR1 |
| 531 | OSR2 |
| 532 | OTP |
| 533 | OTX1 |
| 534 | OTX2 |
| 535 | OVOL1 |
| 536 | OVOL2 |
| 537 | PATZ1 |
| 538 | PAX1 |
| 539 | PAX2 |
| 540 | PAX3 |
| 541 | PAX4 |
| 542 | PAX5 |
| 543 | PAX6 |
| 544 | PAX7 |
| 545 | PAX8 |
| 546 | PAX9 |
| 547 | PBX1 |
| 548 | PBX2 |
| 549 | PBX3 |
| 550 | PBX4 |
| 551 | PDX1 |
| 552 | PGR |
| 553 | PHOX2A |
| 554 | PHOX2B |
| 555 | PITX1 |
| 556 | PITX2 |
| 557 | PITX3 |
| 558 | PKNOX1 |
| 559 | PKNOX2 |
| 560 | PLAG1 |
| 561 | PLAGL1 |
| 562 | POU1F1 |
| 563 | POU2F1 |
| 564 | POU2F2 |
| 565 | POU2F3 |
| 566 | POU3F1 |
| 567 | POU3F2 |
| 568 | POU3F3 |
| 569 | POU3F4 |
| 570 | POU4F1 |
| 571 | POU4F2 |
| 572 | POU4F3 |
| 573 | POU5F1 |
| 574 | POU5F1B |
| 575 | POU6F1 |
| 576 | POU6F2 |
| 577 | PPARA |
| 578 | PPARD |
| 579 | PPARG |
| 580 | PRDM1 |
| 581 | PRDM14 |
| 582 | PRDM16 |
| 583 | PRDM4 |
| 584 | PRDM6 |
| 585 | PRDM9 |
| 586 | PROP1 |
| 587 | PROX1 |
| 588 | PROX2 |
| 589 | PRRX1 |
| 590 | PRRX2 |
| 591 | PTF1A |
| 592 | PURA |
| 593 | RARA |
| 594 | RARB |
| 595 | RARG |
| 596 | RAX |
| 597 | RAX2 |
| 598 | RBAK |
| 599 | RBPJ |
| 600 | RBPJL |
| 601 | REL |
| 602 | RELA |
| 603 | RELB |
| 604 | REST |
| 605 | RFX2 |
| 606 | RFX3 |
| 607 | RFX4 |
| 608 | RFX5 |
| 609 | RFX6 |
| 610 | RFX7 |
| 611 | RHOXF1 |
| 612 | RHOXF2 |
| 613 | RHOXF2B |
| 614 | RORA |
| 615 | RORB |
| 616 | RORC |
| 617 | RREB1 |
| 618 | RUNX1 |
| 619 | RUNX2 |
| 620 | RUNX3 |
| 621 | RXRA |
| 622 | RXRB |
| 623 | RXRG |
| 624 | SCRT1 |
| 625 | SCRT2 |
| 626 | SCX |
| 627 | SEBOX |
| 628 | SHOX |
| 629 | SHOX2 |
| 630 | SIM1 |
| 631 | SIM2 |
| 632 | SIX1 |
| 633 | SIX2 |
| 634 | SIX3 |
| 635 | SIX4 |
| 636 | SIX5 |
| 637 | SIX6 |
| 638 | SKOR1 |
| 639 | SKOR2 |
| 640 | SMAD1 |
| 641 | SMAD3 |
| 642 | SMAD4 |
| 643 | SMAD5 |
| 644 | SMAD9 |
| 645 | SNAI1 |
| 646 | SNAI2 |
| 647 | SNAI3 |
| 648 | SOHLH2 |
| 649 | SOX1 |
| 650 | SOX10 |
| 651 | SOX11 |
| 652 | SOX12 |
| 653 | SOX13 |
| 654 | SOX14 |
| 655 | SOX15 |
| 656 | SOX17 |
| 657 | SOX18 |
| 658 | SOX2 |
| 659 | SOX21 |
| 660 | SOX3 |
| 661 | SOX30 |
| 662 | SOX4 |
| 663 | SOX5 |
| 664 | SOX6 |
| 665 | SOX7 |
| 666 | SOX8 |
| 667 | SOX9 |
| 668 | SP1 |
| 669 | SP2 |
| 670 | SP3 |
| 671 | SP4 |
| 672 | SP5 |
| 673 | SP6 |
| 674 | SP7 |
| 675 | SP8 |
| 676 | SP9 |
| 677 | SPDEF |
| 678 | SPI1 |
| 679 | SPIB |
| 680 | SPIC |
| 681 | SPZ1 |
| 682 | SREBF1 |
| 683 | SREBF2 |
| 684 | SRF |
| 685 | ST18 |
| 686 | STAT1 |
| 687 | STAT2 |
| 688 | STAT3 |
| 689 | STAT4 |
| 690 | STAT5A |
| 691 | STAT5B |
| 692 | STAT6 |
| 693 | T |
| 694 | TAL1 |
| 695 | TAL2 |
| 696 | TBP |
| 697 | TBPL2 |
| 698 | TBR1 |
| 699 | TBX1 |
| 700 | TBX10 |
| 701 | TBX15 |
| 702 | TBX18 |
| 703 | TBX19 |
| 704 | TBX2 |
| 705 | TBX20 |
| 706 | TBX21 |
| 707 | TBX22 |
| 708 | TBX3 |
| 709 | TBX4 |
| 710 | TBX5 |
| 711 | TBX6 |
| 712 | TCF12 |
| 713 | TCF15 |
| 714 | TCF21 |
| 715 | TCF23 |
| 716 | TCF24 |
| 717 | TCF3 |
| 718 | TCF4 |
| 719 | TCF7 |
| 720 | TCF7L1 |
| 721 | TCF7L2 |
| 722 | TCFL5 |
| 723 | TEAD1 |
| 724 | TEAD2 |
| 725 | TEAD3 |
| 726 | TEAD4 |
| 727 | TEF |
| 728 | TERF2 |
| 729 | TET1 |
| 730 | TFAP2A |
| 731 | TFAP2B |
| 732 | TFAP2C |
| 733 | TFAP2D |
| 734 | TFAP2E |
| 735 | TFAP4 |
| 736 | TFCP2 |
| 737 | TFCP2L1 |
| 738 | TFDP1 |
| 739 | TFDP3 |
| 740 | TFE3 |
| 741 | TFEB |
| 742 | TFEC |
| 743 | TGIF1 |
| 744 | TGIF2 |
| 745 | TGIF2LX |
| 746 | THAP1 |
| 747 | THRA |
| 748 | THRB |
| 749 | TLX1 |
| 750 | TLX2 |
| 751 | TLX3 |
| 752 | TOPORS |
| 753 | TP53 |
| 754 | TP63 |
| 755 | TP73 |
| 756 | TWIST1 |
| 757 | TWIST2 |
| 758 | UBP1 |
| 759 | UNCX |
| 760 | USF1 |
| 761 | USF2 |
| 762 | VAX1 |
| 763 | VAX2 |
| 764 | VDR |
| 765 | VENTX |
| 766 | VSX1 |
| 767 | VSX2 |
| 768 | WT1 |
| 769 | XBP1 |
| 770 | XPA |
| 771 | YBX1 |
| 772 | YBX2 |
| 773 | YBX3 |
| 774 | YY1 |
| 775 | YY2 |
| 776 | ZBED1 |
| 777 | ZBTB1 |
| 778 | ZBTB12 |
| 779 | ZBTB14 |
| 780 | ZBTB18 |
| 781 | ZBTB2 |
| 782 | ZBTB20 |
| 783 | ZBTB22 |
| 784 | ZBTB26 |
| 785 | ZBTB3 |
| 786 | ZBTB32 |
| 787 | ZBTB33 |
| 788 | ZBTB34 |
| 789 | ZBTB37 |
| 790 | ZBTB4 |
| 791 | ZBTB42 |
| 792 | ZBTB43 |
| 793 | ZBTB45 |
| 794 | ZBTB48 |
| 795 | ZBTB49 |
| 796 | ZBTB6 |
| 797 | ZBTB7A |
| 798 | ZBTB7B |
| 799 | ZBTB7C |
| 800 | ZEB1 |
| 801 | ZEB2 |
| 802 | ZFHX2 |
| 803 | ZFHX3 |
| 804 | ZFHX4 |
| 805 | ZFP1 |
| 806 | ZFP14 |
| 807 | ZFP2 |
| 808 | ZFP28 |
| 809 | ZFP3 |
| 810 | ZFP30 |
| 811 | ZFP42 |
| 812 | ZFP57 |
| 813 | ZFP64 |
| 814 | ZFP69 |
| 815 | ZFP69B |
| 816 | ZFP82 |
| 817 | ZFP90 |
| 818 | ZFX |
| 819 | ZHX1 |
| 820 | ZIC1 |
| 821 | ZIC2 |
| 822 | ZIC3 |
| 823 | ZIC4 |
| 824 | ZIC5 |
| 825 | ZIK1 |
| 826 | ZIM2 |
| 827 | ZIM3 |
| 828 | ZKSCAN1 |
| 829 | ZKSCAN2 |
| 830 | ZKSCAN3 |
| 831 | ZKSCAN5 |
| 832 | ZKSCAN7 |
| 833 | ZNF10 |
| 834 | ZNF100 |
| 835 | ZNF101 |
| 836 | ZNF114 |
| 837 | ZNF12 |
| 838 | ZNF121 |
| 839 | ZNF124 |
| 840 | ZNF132 |
| 841 | ZNF133 |
| 842 | ZNF134 |
| 843 | ZNF135 |
| 844 | ZNF136 |
| 845 | ZNF140 |
| 846 | ZNF141 |
| 847 | ZNF143 |
| 848 | ZNF146 |
| 849 | ZNF148 |
| 850 | ZNF154 |
| 851 | ZNF157 |
| 852 | ZNF16 |
| 853 | ZNF169 |
| 854 | ZNF17 |
| 855 | ZNF174 |
| 856 | ZNF175 |
| 857 | ZNF177 |
| 858 | ZNF18 |
| 859 | ZNF180 |
| 860 | ZNF181 |
| 861 | ZNF182 |
| 862 | ZNF184 |
| 863 | ZNF189 |
| 864 | ZNF19 |
| 865 | ZNF197 |
| 866 | ZNF2 |
| 867 | ZNF200 |
| 868 | ZNF202 |
| 869 | ZNF205 |
| 870 | ZNF211 |
| 871 | ZNF212 |
| 872 | ZNF213 |
| 873 | ZNF214 |
| 874 | ZNF219 |
| 875 | ZNF22 |
| 876 | ZNF222 |
| 877 | ZNF223 |
| 878 | ZNF224 |
| 879 | ZNF225 |
| 880 | ZNF232 |
| 881 | ZNF235 |
| 882 | ZNF248 |
| 883 | ZNF25 |
| 884 | ZNF250 |
| 885 | ZNF254 |
| 886 | ZNF257 |
| 887 | ZNF26 |
| 888 | ZNF260 |
| 889 | ZNF263 |
| 890 | ZNF264 |
| 891 | ZNF266 |
| 892 | ZNF267 |
| 893 | ZNF273 |
| 894 | ZNF274 |
| 895 | ZNF276 |
| 896 | ZNF28 |
| 897 | ZNF280A |
| 898 | ZNF281 |
| 899 | ZNF282 |
| 900 | ZNF283 |
| 901 | ZNF284 |
| 902 | ZNF285 |
| 903 | ZNF287 |
| 904 | ZNF296 |
| 905 | ZNF3 |
| 906 | ZNF30 |
| 907 | ZNF300 |
| 908 | ZNF302 |
| 909 | ZNF304 |
| 910 | ZNF311 |
| 911 | ZNF317 |
| 912 | ZNF32 |
| 913 | ZNF320 |
| 914 | ZNF322 |
| 915 | ZNF324 |
| 916 | ZNF324B |
| 917 | ZNF329 |
| 918 | ZNF331 |
| 919 | ZNF333 |
| 920 | ZNF334 |
| 921 | ZNF337 |
| 922 | ZNF33A |
| 923 | ZNF33B |
| 924 | ZNF34 |
| 925 | ZNF341 |
| 926 | ZNF343 |
| 927 | ZNF345 |
| 928 | ZNF35 |
| 929 | ZNF350 |
| 930 | ZNF354A |
| 931 | ZNF354B |
| 932 | ZNF354C |
| 933 | ZNF37A |
| 934 | ZNF382 |
| 935 | ZNF383 |
| 936 | ZNF384 |
| 937 | ZNF385D |
| 938 | ZNF394 |
| 939 | ZNF396 |
| 940 | ZNF398 |
| 941 | ZNF41 |
| 942 | ZNF410 |
| 943 | ZNF415 |
| 944 | ZNF416 |
| 945 | ZNF417 |
| 946 | ZNF418 |
| 947 | ZNF419 |
| 948 | ZNF423 |
| 949 | ZNF425 |
| 950 | ZNF429 |
| 951 | ZNF430 |
| 952 | ZNF431 |
| 953 | ZNF432 |
| 954 | ZNF433 |
| 955 | ZNF436 |
| 956 | ZNF439 |
| 957 | ZNF44 |
| 958 | ZNF440 |
| 959 | ZNF441 |
| 960 | ZNF442 |
| 961 | ZNF443 |
| 962 | ZNF444 |
| 963 | ZNF445 |
| 964 | ZNF449 |
| 965 | ZNF45 |
| 966 | ZNF454 |
| 967 | ZNF460 |
| 968 | ZNF467 |
| 969 | ZNF468 |
| 970 | ZNF479 |
| 971 | ZNF480 |
| 972 | ZNF483 |
| 973 | ZNF484 |
| 974 | ZNF485 |
| 975 | ZNF486 |
| 976 | ZNF487 |
| 977 | ZNF490 |
| 978 | ZNF492 |
| 979 | ZNF496 |
| 980 | ZNF501 |
| 981 | ZNF502 |
| 982 | ZNF506 |
| 983 | ZNF513 |
| 984 | ZNF519 |
| 985 | ZNF524 |
| 986 | ZNF525 |
| 987 | ZNF527 |
| 988 | ZNF528 |
| 989 | ZNF529 |
| 990 | ZNF530 |
| 991 | ZNF534 |
| 992 | ZNF540 |
| 993 | ZNF543 |
| 994 | ZNF547 |
| 995 | ZNF548 |
| 996 | ZNF549 |
| 997 | ZNF550 |
| 998 | ZNF552 |
| 999 | ZNF554 |
| 1000 | ZNF555 |
| 1001 | ZNF557 |
| 1002 | ZNF558 |
| 1003 | ZNF561 |
| 1004 | ZNF562 |
| 1005 | ZNF563 |
| 1006 | ZNF564 |
| 1007 | ZNF565 |
| 1008 | ZNF566 |
| 1009 | ZNF567 |
| 1010 | ZNF570 |
| 1011 | ZNF571 |
| 1012 | ZNF573 |
| 1013 | ZNF574 |
| 1014 | ZNF580 |
| 1015 | ZNF582 |
| 1016 | ZNF584 |
| 1017 | ZNF585A |
| 1018 | ZNF586 |
| 1019 | ZNF587 |
| 1020 | ZNF589 |
| 1021 | ZNF594 |
| 1022 | ZNF595 |
| 1023 | ZNF596 |
| 1024 | ZNF597 |
| 1025 | ZNF605 |
| 1026 | ZNF610 |
| 1027 | ZNF611 |
| 1028 | ZNF613 |
| 1029 | ZNF614 |
| 1030 | ZNF615 |
| 1031 | ZNF616 |
| 1032 | ZNF619 |
| 1033 | ZNF620 |
| 1034 | ZNF621 |
| 1035 | ZNF626 |
| 1036 | ZNF627 |
| 1037 | ZNF641 |
| 1038 | ZNF649 |
| 1039 | ZNF653 |
| 1040 | ZNF655 |
| 1041 | ZNF660 |
| 1042 | ZNF662 |
| 1043 | ZNF667 |
| 1044 | ZNF669 |
| 1045 | ZNF671 |
| 1046 | ZNF674 |
| 1047 | ZNF675 |
| 1048 | ZNF677 |
| 1049 | ZNF680 |
| 1050 | ZNF681 |
| 1051 | ZNF682 |
| 1052 | ZNF684 |
| 1053 | ZNF69 |
| 1054 | ZNF691 |
| 1055 | ZNF692 |
| 1056 | ZNF695 |
| 1057 | ZNF7 |
| 1058 | ZNF701 |
| 1059 | ZNF704 |
| 1060 | ZNF705G |
| 1061 | ZNF707 |
| 1062 | ZNF708 |
| 1063 | ZNF71 |
| 1064 | ZNF711 |
| 1065 | ZNF713 |
| 1066 | ZNF714 |
| 1067 | ZNF716 |
| 1068 | ZNF730 |
| 1069 | ZNF736 |
| 1070 | ZNF737 |
| 1071 | ZNF74 |
| 1072 | ZNF740 |
| 1073 | ZNF749 |
| 1074 | ZNF75A |
| 1075 | ZNF75D |
| 1076 | ZNF76 |
| 1077 | ZNF764 |
| 1078 | ZNF765 |
| 1079 | ZNF766 |
| 1080 | ZNF768 |
| 1081 | ZNF77 |
| 1082 | ZNF770 |
| 1083 | ZNF771 |
| 1084 | ZNF774 |
| 1085 | ZNF776 |
| 1086 | ZNF777 |
| 1087 | ZNF778 |
| 1088 | ZNF780A |
| 1089 | ZNF782 |
| 1090 | ZNF783 |
| 1091 | ZNF784 |
| 1092 | ZNF785 |
| 1093 | ZNF786 |
| 1094 | ZNF787 |
| 1095 | ZNF789 |
| 1096 | ZNF79 |
| 1097 | ZNF790 |
| 1098 | ZNF791 |
| 1099 | ZNF792 |
| 1100 | ZNF793 |
| 1101 | ZNF799 |
| 1102 | ZNF8 |
| 1103 | ZNF805 |
| 1104 | ZNF808 |
| 1105 | ZNF81 |
| 1106 | ZNF816 |
| 1107 | ZNF821 |
| 1108 | ZNF823 |
| 1109 | ZNF84 |
| 1110 | ZNF846 |
| 1111 | ZNF85 |
| 1112 | ZNF852 |
| 1113 | ZNF860 |
| 1114 | ZNF879 |
| 1115 | ZNF880 |
| 1116 | ZNF891 |
| 1117 | ZNF90 |
| 1118 | ZNF93 |
| 1119 | ZNF98 |
| 1120 | ZSCAN1 |
| 1121 | ZSCAN10 |
| 1122 | ZSCAN16 |
| 1123 | ZSCAN22 |
| 1124 | ZSCAN23 |
| 1125 | ZSCAN26 |
| 1126 | ZSCAN29 |
| 1127 | ZSCAN30 |
| 1128 | ZSCAN31 |
| 1129 | ZSCAN4 |
| 1130 | ZSCAN5A |
| 1131 | ZSCAN5C |
| 1132 | ZSCAN9 |
